# Natural selection interacts with the local recombination rate to shape the evolution of hybrid genomes

**DOI:** 10.1101/212407

**Authors:** Molly Schumer, Chenling Xu, Daniel L. Powell, Arun Durvasula, Laurits Skov, Chris Holland, Sriram Sankararaman, Peter Andolfatto, Gil G. Rosenthal, Molly Przeworski

**Affiliations:** Hanna H. Gray Fellow, Howard Hughes Medical Institute; Harvard Society of Fellows, Harvard University; Department of Biological Sciences, Columbia University; Centro de Investigaciones Científicas de las Huastecas “Aguazarca”; Center for Computational Biology, University of California at Berkeley; Department of Biology, Texas A&M University; Department of Human Genetics, David Geffen School of Medicine, University of California, Los Angeles; Bioinformatics Research Centre, Aarhus University, Aarhus C., Denmark; Department of Computer Science, University of California, Los Angeles; Department of Ecology and Evolutionary Biology and Lewis-Sigler Institute for Integrative Genomics, Princeton University; Department of Systems Biology, Columbia University

## Abstract

While hybridization between species is increasingly appreciated to be a common occurrence, little is known about the forces that govern the subsequent evolution of hybrid genomes. We considered this question in three independent, naturally-occurring hybrid populations formed between swordtail fish species *Xiphophorus birchmanni* and *X. malinche.* To this end, we built a fine-scale genetic map and inferred patterns of local ancestry along the genomes of 690 individuals sampled from the three populations. In all three cases, we found hybrid ancestry to be more common in regions of high recombination and where there is linkage to fewer putative targets of selection. These same patterns are also apparent in a reanalysis of human-Neanderthal admixture. Our results lend support to models in which ancestry from the “minor” parental species persists only where it is rapidly uncoupled from alleles that are deleterious in hybrids, and show the retention of hybrid ancestry to be at least in part predictable from genomic features. Our analyses further indicate that in swordtail fish, the dominant source of selection on hybrids stems from deleterious combinations of epistatically-interacting alleles.

**One sentence summary:** The persistence of hybrid ancestry is predictable from local recombination rates, in three replicate hybrid populations as well as in humans.

## Main text

Understanding speciation is central to understanding evolution, but so much about the process still puzzles us. The foundational work in evolutionary biology envisioned speciation as an ordered process during which reproductive barriers, once established, prevent gene flow between species (*1*). We now realize, however, that speciation is much more dynamic, with hybridization occurring both during and after the evolution of reproductive barriers and evidence of past hybridization with close relatives still visible in the genomes of myriad animal and plant species (*2–9*). The ubiquity of hybridization raises the question of how species that hybridize remain genetically and ecologically differentiated.

At least part of the answer is likely that selection filters out deleterious hybrid ancestry from the genome (*1*). For instance, in hominins and swordtail fish, individuals are less likely to carry hybrid ancestry near functionally important elements (*6, 10, 11*), presumably because it is especially deleterious in such regions. Aside from these observations, however, little is known about how admixed genomes evolve—or, in some cases, stabilize—following hybridization. Our understanding of the evolution of hybrid genomes is complicated by the existence of many possible modes of selection and by the fact that, in most systems, the location of sites under selection is unknown. Decades of experimental work have demonstrated that Dobzhansky-Muller incompatibilities (DMIs) are a central mechanism underlying reproductive isolation once species are formed (*12–16*), but the importance of DMIs in shaping the evolution of hybrid genomes remains unknown, as does the role of other modes of selection. Notably, it was recently pointed out that when there is introgression from a species with a lower effective population size, hybrids may suffer from increased genetic load (“hybridization load”) due to the introduction of weakly deleterious alleles (*10, 17*). Depending on the environment in which hybrids find themselves, alleles that underlie ecological adaptations in the parental species may also be deleterious in hybrids (*18, 19*). Complicating matters yet further, modes of selection on hybrid ancestry will likely vary from system to system, depending on the extent of genetic divergence and ecological differentiation between the parental species, as well as long-term differences in their effective population sizes.

One feature, however, is expected to play a central role in all these models: variation in recombination rates along the genome (*10, 17, 20–22*). Theory predicts that selection is more likely to weed out hybrid ancestry in regions of low recombination (*23–25*). Specifically, in models of DMIs, minor parent ancestry will persist preferentially in regions of higher recombination because it is more rapidly uncoupled from mutations that are incompatible with the prevalent (i.e., major parent) genetic background (Fig. 1). Similarly, in models of hybridization load, all else being equal, shorter linkage blocks will carry fewer weakly deleterious mutations and therefore be will be less rapidly purged by selection (*10, 17*; Fig. S1). Previous studies have reported patterns potentially consistent with these expectations (*26, 27*), but without directly investigating ancestry patterns and their relationship with local recombination rates (*28*).

**Figure 1.**
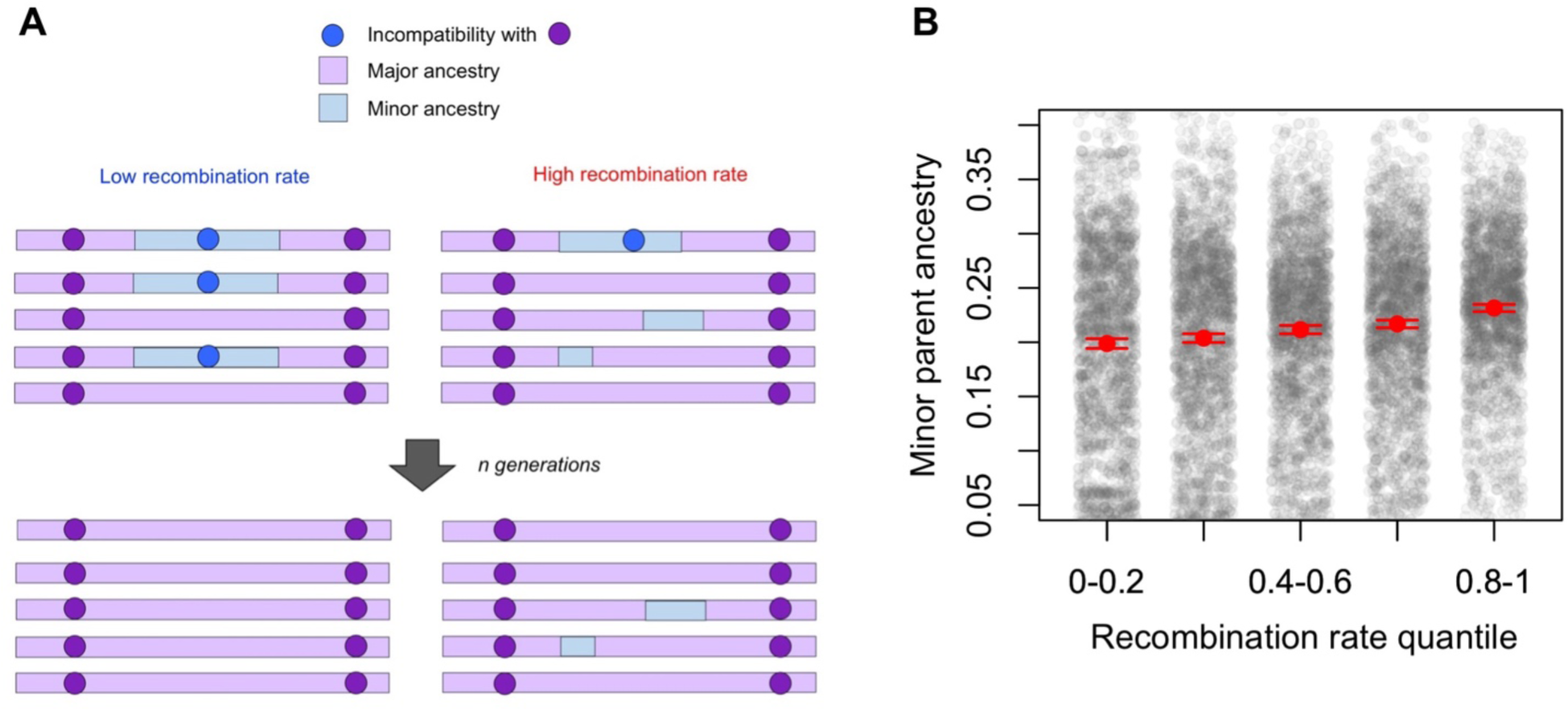
Predicted relationship between minor parent ancestry and recombination rates in the presence of hybrid incompatibilities. (A) In the presence of hybrid incompatibilities, the local recombination rate influences the rate at which neutral regions will be uncoupled from nearby hybrid incompatibility loci; as a result, more minor parent ancestry should be retained in regions of high recombination. (B) This prediction is borne out in simulations. Shown here are simulation results with two pairs of randomly placed hybrid incompatibilities per chromosome and *s*=0.1, plausible parameters for *Xiphophorus* species (*29*, Supporting Information 5). In these simulations, 250 individuals were sampled at generation 70 and ancestry along 25 Mb chromosomes was summarized in 50 kb windows. Shown here is a replicate selected at random, with the number of simulated chromosomes chosen to mimic the amount of data used in our analyses; 74% of simulations had a significantly positive relationship between minor parent ancestry and recombination rate at the 5% level. Red points and whiskers indicate the mean minor parent ancestry with two standard errors of the mean determined by bootstrapping windows; gray points show raw data. Note that the y-axis range is truncated.

### Recombination shapes ancestry in swordtail fish

To test predictions about the role of recombination in filtering hybrid ancestry, we took advantage of a set of naturally occurring hybrid populations between two swordtail fish species, *Xiphophorus birchmanni* and a closely related species, *X. malinche* (Fig. 2; Supporting Information 1-3). The two species are ~0.5% divergent at the nucleotide level and incomplete lineage sorting between the two is relatively rare (*29*; Fig. 2A). We focused on three hybrid populations that formed independently between the two fewer than 100 generations ago (*29*), likely as a result of human-mediated habitat disturbance (*30*). Previous analyses of hybrid zones between these two species, including two of the three populations analyzed here, suggested that there are on the order of 100 pairs of unlinked DMIs segregating in hybrids (*29, 31*), with estimated selection coefficients ~0.03-0.05 (*29*), and potentially many more linked DMIs, indicating that swordtail hybrids may be experiencing widespread selection on DMIs.

**Figure 2.**
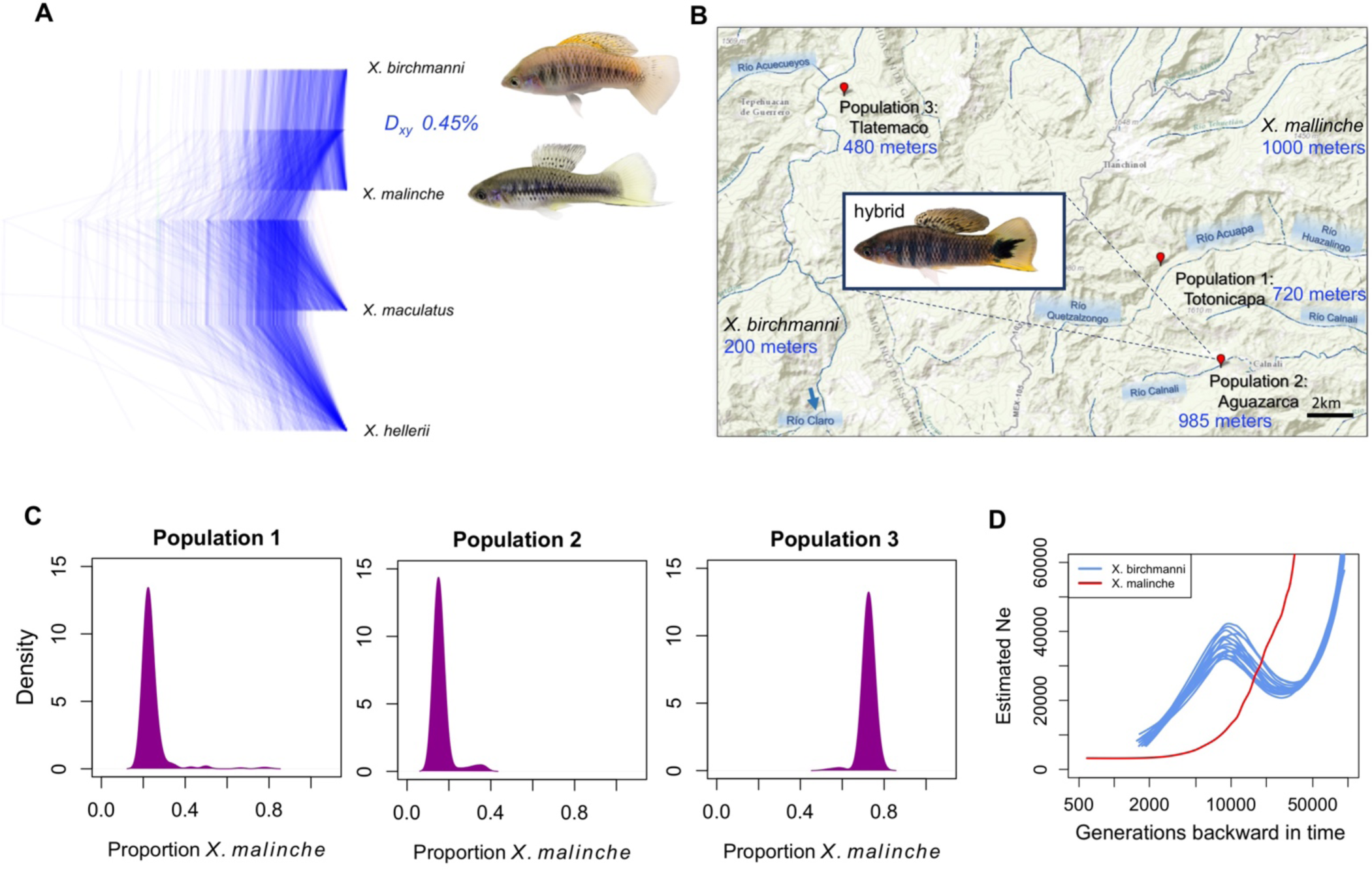
Hybridization between sister species *X. birchmanni* and *X. malinche*. (A) Maximum likelihood trees produced by RAxML (*54*) for 1,000 alignments of randomly selected 10 kb regions. D_xy_ refers to the average nucleotide divergence between *X. birchmanni* and *X. malinche*. Outgroup genome sequences were obtained from previous work (*55, 56*). (B) Locations of the three hybrid populations on which we focused, all sampled from different river systems in Hidalgo, Mexico; listed in blue are elevations of the sampled hybrid populations and typical elevations for the parental populations. (C) Inferred ancestry proportions of individuals from each of the three hybrid populations (see Methods for details). (D) Loess fit to effective population size estimates inferred by MSMC from one *X. malinche* genome and each of the 20 *X. birchmanni* genomes collected for this study, assuming a mutation rate of 3.5 × 10^−9^ (*57*; see Supporting Information 3). The time interval overlapping with zero is not plotted.

To infer local ancestry patterns, we generated ~1X low coverage whole genome data for 690 hybrids sampled from the three hybrid populations (Supporting Information 1). We estimated local ancestry patterns for the 690 hybrids by applying a hidden Markov model (*32*); this approach is predicted to have high accuracy for these hybrid populations, given the marker density and time since mixture (*29, 32, 33*). Using ancestry calls at 1-1.2 million sites genome wide, we inferred that two of the hybrid populations derive on average 75-80% of their genomes from *X. birchmanni*, whereas individuals in the third population derive on average 72% of their genomes from *X. malinche* (Fig. 2; Supporting Information 1; *34*). The median homozygous tract length for the minor parent ranges from 84 kb to 225 kb across the three populations, roughly matching expectations for hybrid populations of these ages and mixture proportions (Supporting Information 4).

To consider the relationship between local ancestry and recombination rates, we inferred a fine-scale genetic map for *X. birchmanni* from patterns of linkage disequilibrium (LD) in unrelated individuals (Table S1; Supporting Information 2, 4-5). Based on our previous work on recombination in this taxon (*35*), we had a strong prior expectation that local recombination rates should be conserved between *X. birchmanni* and *X. malinche* (Supporting Information 6). We also generated crossover maps from hybrids based on inferred switch points between the two ancestries. Overall the hybrid and parental maps are consistent (Fig. S2), with the correlations between maps roughly comparable to what would be expected if the maps were in fact identical (Supporting Information 7).

In all three hybrid populations of swordtail fish, the probability of carrying ancestry from the minor parent increases with the local recombination rate (Fig. 3, Table 1). This association remains irrespective of the choice of scale (Fig. S3) and after thinning the SNP and ancestry variation data to control for possible differences in the ability to reliably infer recombination rates or the power to call hybrid ancestry across windows (Supporting Information 4). The preferential persistence of minor parent ancestry in regions of higher recombination is not expected under neutrality (Fig. S1) and instead indicates that minor parent ancestry was retained where it was more likely to have been rapidly uncoupled from the deleterious alleles with which it was originally linked (Supporting Information 5). This qualitative pattern can be generated under several models of selection, including selection against DMIs, selection against weakly deleterious alleles introduced by hybridization, or widespread ecological selection against loci that derive from the minor parent (Fig. 1, Fig. S1).

**Figure 3.**
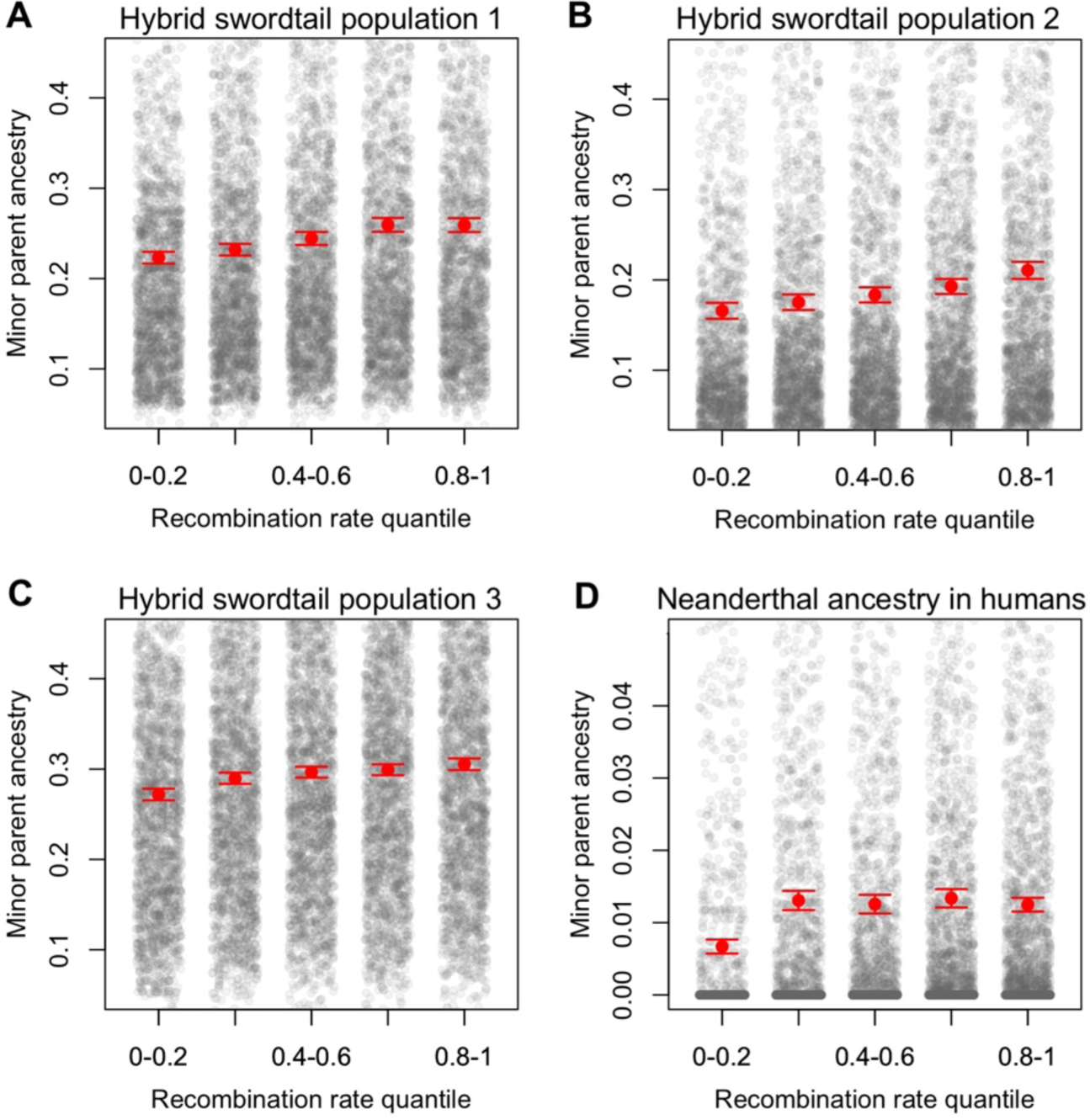
Minor parent ancestry is significantly decreased in regions of low recombination, in swordtail fish and humans. (A-C) Minor parent ancestry is rarest in regions of the swordtail genome with the lowest recombination rates. (D) Neanderthal ancestry is likewise rarest in regions of the human genome with the lowest recombination rates. Removing windows of particularly high Neanderthal ancestry, which may have experienced adaptive introgression, further strengthens this relationship (Fig. S8). Data are summarized in 50 kb windows in swordtail analyses and 250 kb windows in analysis of human data; results are similar for a range of window sizes (summarized in Table 1). Quantile binning is for visual representation only; all statistical tests reported in Table 1 were performed on the unbinned data. Red points and whiskers indicate the mean minor parent ancestry and two standard errors of the mean, obtained by bootstrapping windows; gray points show raw data. Note that the y-axis is truncated in all panels.

**Table 1.** Relationship between minor parent ancestry and recombination rate. Results for Spearman’s correlations between minor parent ancestry and recombination rate at several scales, in three swordtail fish hybrid populations and for archaic hominin ancestry in the human genome. Also shown are partial correlation results, controlling for the number of coding base pairs. Because different filtering approaches were applied to the archaic hominin ancestry datasets in the original studies, results reported here are based on windows included in all datasets. P-values are obtained after thinning windows, to minimize correlations among nearby windows (see Supporting Information 4). Additional information and analyses can be found in Supporting Information 4 and Supporting Information 9.

| Population | Spearman's correlation<br>between minor ancestry and rate |  |  | Spearman's partial correlation<br>between minor ancestry and rate<br>including coding base pairs |  |  |
| --- | --- | --- | --- | --- | --- | --- |
|  | 50 kb | 250 kb | 500 kb | 50 kb | 250 kb | 500 kb |
| **Swordtail population 1** | $\rho = 0.14$<br>$p = 10^{-10}$ | $\rho = 0.15$<br>$p = 10^{-8}$ | $\rho = 0.18$<br>$p = 10^{-10}$ | $\rho = 0.15$<br>$p = 10^{-11}$ | $\rho = 0.16$<br>$p = 10^{-8}$ | $\rho = 0.18$<br>$p = 10^{-10}$ |
| **Swordtail population 2** | $\rho = 0.1$<br>$p = 10^{-4}$ | $\rho = 0.10$<br>$p = 10^{-4}$ | $\rho = 0.08$<br>$p = 0.007$ | $\rho = 0.1$<br>$p = 10^{-4}$ | $\rho = 0.11$<br>$p = 10^{-4}$ | $\rho = 0.08$<br>$p = 0.008$ |
| **Swordtail population 3** | $\rho = 0.11$<br>$p = 10^{-6}$ | $\rho = 0.11$<br>$p = 10^{-4}$ | $\rho = 0.12$<br>$p = 10^{-5}$ | $\rho = 0.11$<br>$p = 10^{-6}$ | $\rho = 0.11$<br>$p = 10^{-4}$ | $\rho = 0.11$<br>$p = 10^{-4}$ |
| **Neanderthal ancestry in humans**<br>(diCal-admix) | $\rho = 0.08$<br>$p = 10^{-14}$ | $\rho = 0.17$<br>$p = 10^{-35}$ | $\rho = 0.19$<br>$p = 10^{-42}$ | $\rho = 0.08$<br>$p = 10^{-13}$ | $\rho = 0.17$<br>$p = 10^{-34}$ | $\rho = 0.19$<br>$p = 10^{-43}$ |
| **Neanderthal ancestry in humans**<br>(Sankararaman et al. 2014) | $\rho = 0.06$<br>$p = 10^{-8}$ | $\rho = 0.15$<br>$p = 10^{-26}$ | $\rho = 0.16$<br>$p = 10^{-34}$ | $\rho = 0.06$<br>$p = 10^{-8}$ | $\rho = 0.14$<br>$p = 10^{-25}$ | $\rho = 0.17$<br>$p = 10^{-34}$ |
| **Neanderthal ancestry in humans**<br>(Skov et al. 2017) | $\rho = 0.07$<br>$p = 10^{-10}$ | $\rho = 0.14$<br>$p = 10^{-24}$ | $\rho = 0.15$<br>$p = 10^{-30}$ | $\rho = 0.07$<br>$p = 10^{-9}$ | $\rho = 0.14$<br>$p = 10^{-24}$ | $\rho = 0.16$<br>$p = 10^{-30}$ |
| **Denisovan ancestry in humans**<br>(Sankararaman et al. 2016) | $\rho = 0.02$<br>$p = 0.06$ | $\rho = 0.05$<br>$p = 10^{-4}$ | $\rho = 0.07$<br>$p = 10^{-6}$ | $\rho = 0.02$<br>$p = 0.07$ | $\rho = 0.05$<br>$p = 10^{-4}$ | $\rho = 0.07$<br>$p = 10^{-6}$ |

In principle, the retention of hybrid ancestry should be most accurately predicted from the exact number of deleterious alleles to which a minor parent segment was linked since hybridization occurred. Local recombination rates are one proxy for this (unknown) parameter, as are the number of coding base pairs nearby. In these data, both factors predict minor parent ancestry (Fig. S4; Fig. S5; Supporting Information 4), but local recombination is a stronger predictor and remains a predictor after controlling for the number of coding base pairs (Table 1; Table S2). These findings are consistent with those obtained in simulations mimicking the data structure (Supporting Information 4-5), presumably because the number of coding base pairs nearby is an extremely noisy proxy.

### The source of selection

Controlling for the recombination rate, local ancestry is positively correlated between all pairs of hybrid populations, with weaker but significant correlations seen even between populations with different major parent ancestries (Fig. 4). These findings are expected from selection on the same underlying loci in independently formed populations (Supporting Information 8). Whereas both DMIs and hybridization load are predicted to drive positive correlations in local ancestry across populations regardless of the admixture proportions (Fig. S6), ecological selection against minor parent ancestry should lead to negative correlations in local ancestry and is thus inconsistent with the observed patterns (Fig. 4; Supporting Information 8).

**Figure 4.**
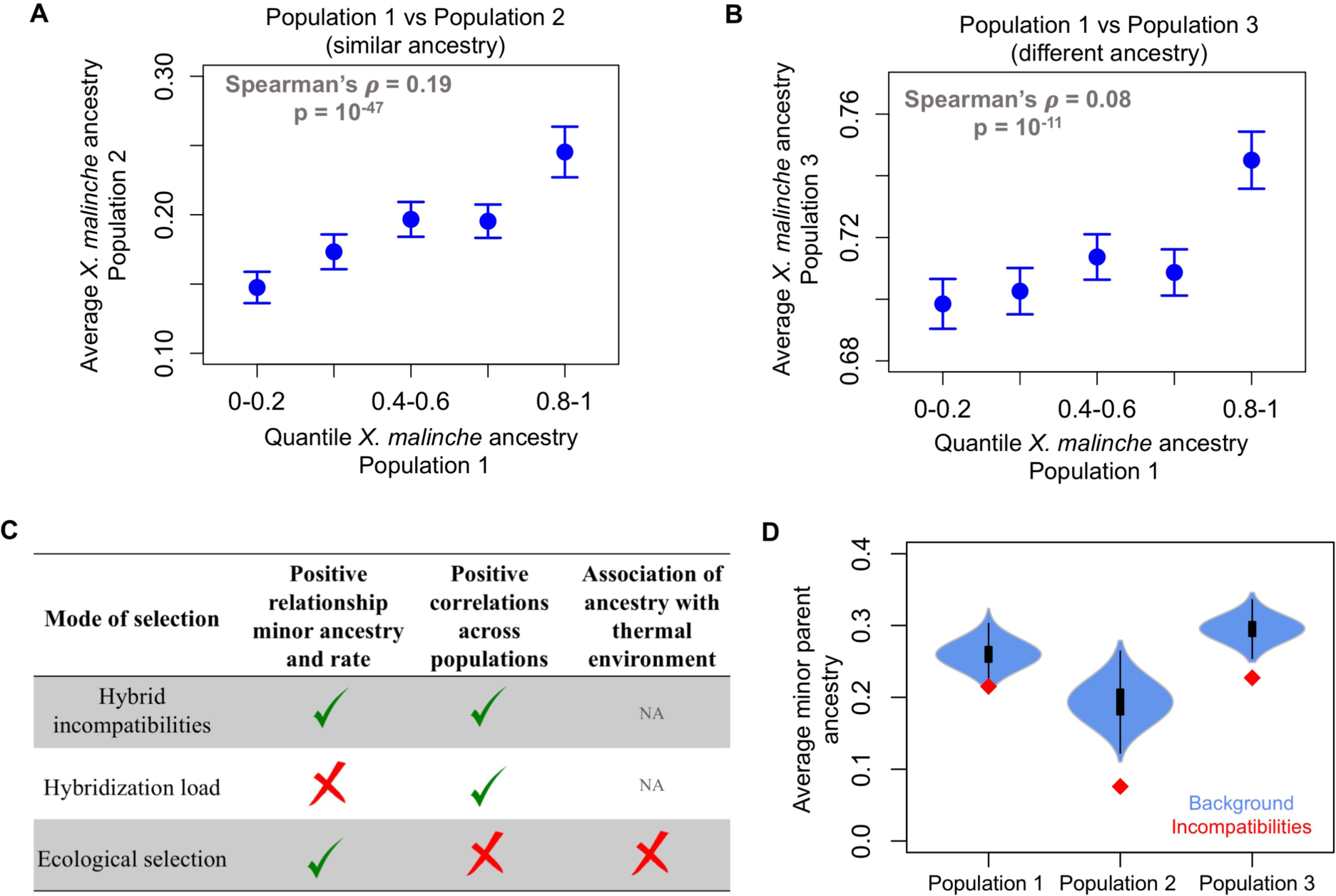
Evidence for DMIs as the major source of selection on hybrids. (A) Local ancestry is strongly and significantly correlated between independently formed swordtail hybrid populations with similar mixture proportions (using 0.1 cM windows; Supporting Information 8). (B) Local ancestry is also correlated between swordtail hybrid populations with distinct major parent ancestry, but the relationship is weaker. Similar results for cross-population correlations in ancestry are observed when controlling for the number of coding base pairs in a window using a partial correlation analysis (when comparing population 1 and 2, ρ = 0.19, p=10^−46^; in turn, ρ 0.04-0.08, p<0.005 for populations 1 and 2 versus population 3). Points show the mean ancestry and whiskers indicate two standard errors of the mean. (C) Predictions for different modes of selection on hybrids. Predictions met in the data are shown with a check and not met with a red cross. See text for details. (D) Average minor parent ancestry is unusually depleted in 50 kb windows containing previously mapped, unlinked DMIs (red points, from *31*) compared to 1,000 null datasets generated by randomly sampling the same number of windows from the background (blue distribution). Simulations suggest that observed ancestry departures at DMIs are not expected as a result of ascertainment (Supporting Information 5).

Comparison among the three hybrid populations also provides a means to distinguish between the remaining two hypotheses. Analyzing genome sequences from *X. malinche* (*5, 29*) and *X. birchmanni,* we found that *X. malinche* has had a lower long-term effective population size than *X. birchmanni* (Fig. 2; Supporting Information 3), as seen both in the approximately four-fold lower average heterozygosity in *X. malinche* (0.03% vs 0.12% per base pair, respectively) and in estimates of effective population sizes over time from high coverage genome sequences (Fig. 2, Table S1). Consistent with a lower long-term effective population size, *X. malinche* carries significantly more putative deleterious alleles relative to the inferred ancestral sequence than does *X. birchmanni*, as measured by the number of derived, non-synonymous substitutions per haploid genome (a 2.5% excess, p=0.016 based on 1,000 bootstrap resamples; see Supporting Information 3; *36, 37*). Because *X. birchmanni* and *X. malinche* source populations differ in the number of putatively deleterious variants, the three hybrid populations of swordtail fish provide an informative contrast: whereas DMIs should lead to selection against minor parent ancestry in all three populations, hybridization load should favor the major parent in populations 1 and 2 and the minor parent in population 3 (Fig. 2; Fig. 4).

In this regard, the fact that minor parent ancestry also increases with recombination in the third hybrid population, which derives most of its genome from the parental species that has lower effective population size (Fig. 2, 3), indicates that hybrid incompatibilities are the dominant mode of selection shaping ancestry in the genome in these hybrid populations, rather than selection against hybridization load (Fig. 4; Fig. S7; Supporting Information 4-5). In principle, ecological selection favoring the major parent could also produce a positive correlation between recombination rate and ancestry (though not the positive correlations in ancestry across populations; Fig. 4). However, this scenario would require two of the hybrid populations to occur in more *birchmanni*-like environments and one in a more *malinche*-like environment, when available evidence suggests otherwise—notably, all of the hybrid populations are found in thermal environments that are mismatched to the environment where their major parent is found (Fig. 2; Supporting Information 5).

Moreover, in all three hybrid populations, minor parent ancestry is unusually low near previously mapped DMIs between the two parental species (*29, 31*), a pattern that should not arise from the approach used to identify DMIs (Supporting Information 5), but is expected from selection on epistatically-interacting alleles (Fig. 4; Supporting Information 4-5). Taken together, these lines of evidence indicate that DMIs are the main (though not necessarily sole) source of selection shaping the retention of hybrid ancestry in these three swordtail fish hybrid populations (Fig. 4).

### Ancestry also interacts with the local recombination rate in hominins

To evaluate the generality of the relationship between recombination rate and ancestry seen in swordtails, we considered the only other case with similar genomic data available: admixture between humans and archaic hominins. Several studies have reported that the average proportion of Neanderthal ancestry decreases with the number of closely linked coding base pairs and with a measure of the strength of purifying selection at linked sites (*6, 10, 17, 38*), patterns for which both DMIs and hybridization load (due to the smaller effective population size of Neanderthals, *39*) have been proposed as explanations (*6, 10, 17*). Reanalyzing the data, we found that the proportion of Neanderthal ancestry (the minor parental species) decreases in regions of the human genome with lower recombination rates (Fig. 3D; Table 1; Table S3). This relationship is seen for different window size choices and with any of three approaches to infer Neanderthal ancestry in the human genome (Table 1), and is not expected as a result of variation in the power to identify introgression along the genome (Supporting Information 9). The effect of local recombination rate on Neanderthal ancestry also persists after accounting for the number of coding base pairs nearby (Table 1; Supporting Information 9). Interestingly, the relationship between Neanderthal ancestry and local recombination rate is especially strong when excluding regions of unusually high frequency Neanderthal ancestry (e.g. top 1%; Fig. S8), possibly because these regions are enriched for cases of adaptive introgression (*6, 38, 40, 41*). Repeating these analyses for Denisovan ancestry, for which there is lower power to identify ancestry tracts, there is a much weaker but consistent trend (Table 1; Supporting Information 9).

As in swordtails, the persistence of Neanderthal ancestry in regions of higher recombination is not expected under neutrality (Fig. S1) but could be generated by selection against DMIs, weakly deleterious alleles introduced by Neanderthals (*6, 10, 17, 38*), or widespread ecological selection against Neanderthal ancestry (Fig. 1, Fig. S1). Unlike in the case of swordtails, however, these causes cannot be distinguished based on these data alone (*6, 10, 17, 38, 42*). Moreover, the conclusion about the source of selection reached for swordtail fish need not hold for hominins, in particular because modern humans and Neanderthals (Denisovans) were less diverged when they are thought to have interbred, and thus may have accumulated many fewer DMIs (*43, 44*).

### The predictability of hybrid ancestry

Hybrid ancestry is predicted by the local recombination rate across three replicate admixture events between the same species pair in swordtail fish, as well as in two cases of admixture in hominins. In swordtail fish hybrids, several lines of evidence indicate that selection against hybrid incompatibilities is the dominant force shaping minor parent ancestry in the genome. In hominins, the source of selection remains unclear. Regardless of the precise mechanisms of selection on hybrids, the generality of these patterns reveals the retention of hybrid ancestry to be at least in part predictable from genomic features.

The relationship of minor parent ancestry to local recombination thus provides a useful tool for predicting where in the genome we might expect hybrid ancestry to persist preferentially. In particular, in hominins, meiotic recombination events are directed to the genome by binding of the *PRDM9* gene, whereas in swordtail fish, they are not and instead are concentrated around CpG islands and other promoter-like features (*35*; Supporting Information 5-6). Accordingly, we found that in swordtail fish, minor parent ancestry is higher around CpG islands and transcription start sites whereas in humans, it is not (Fig. 5; Supporting Information 5). In other words, the mechanism by which recombination is directed to the genome shapes the retention of hybrid ancestry.

**Figure 5.**
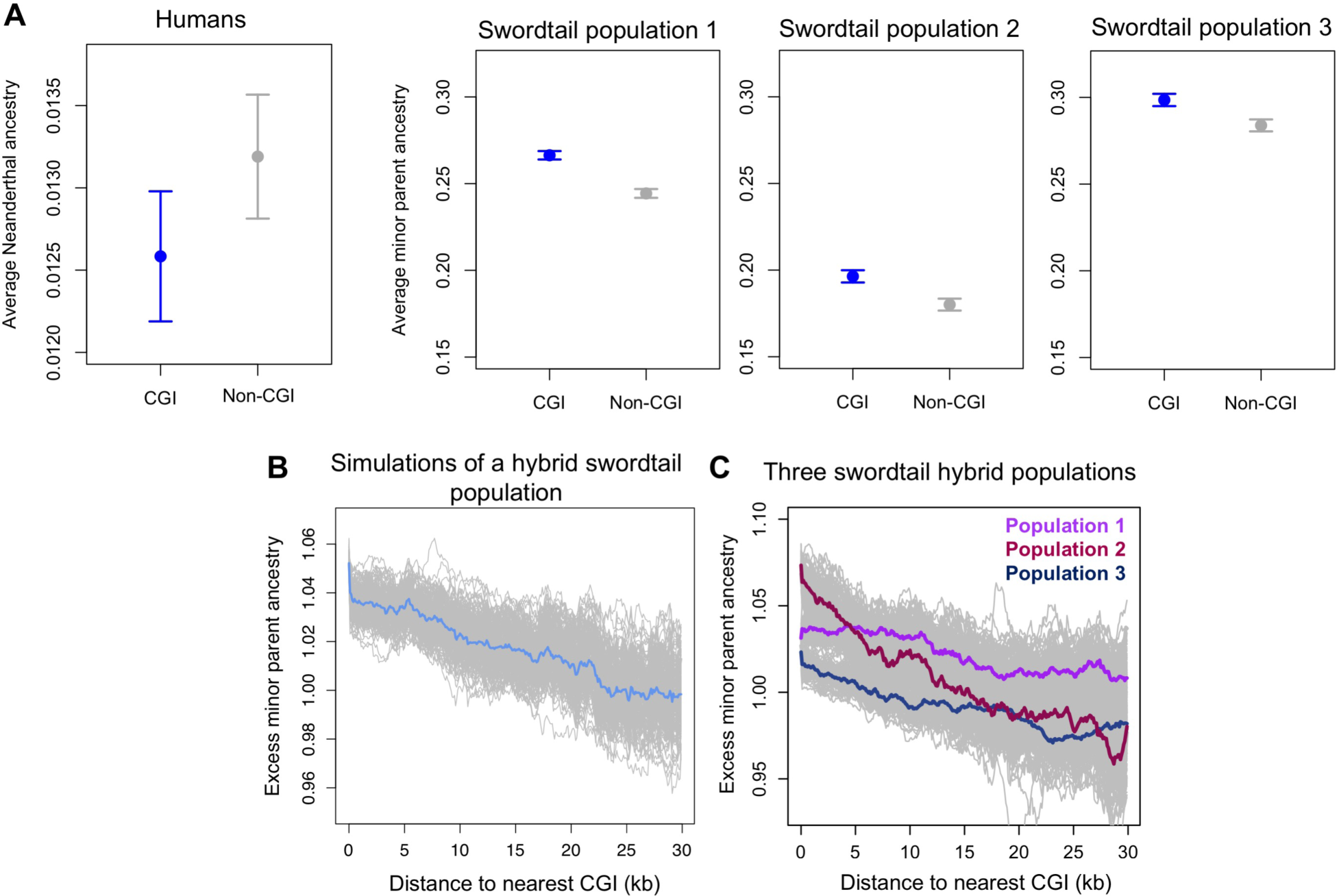
The mechanism by which recombination is directed to the genome shapes the genomic location of minor parent ancestry. (A) Neanderthal ancestry in the human genome is not significantly elevated in 50 kb windows that overlap with CpG islands (CGIs), when compared to windows that do not overlap CGIs but have similar GC base pair composition (the fold difference is 0.95, p=0.91). Points show the mean of each group and whiskers indicate one standard error of the mean obtained by 1,000 joint bootstraps resampling the data. (A) In contrast, in all three swordtail fish populations, minor parent ancestry is significantly elevated in windows that overlap CGIs compared to windows that do not but have similar GC base pair composition (in population 1, the fold-difference is 1.09, p<0.005; in population 2, the fold-difference is 1.09, p<0.005; and in population 3 the fold-difference is 1.02, p<0.005). See Supporting Information 5 for details. (B) Simulations of incompatibility selection in swordtails predict high minor parent ancestry near CGIs (Supporting Information 5). (C) This prediction is met for all three hybrid populations. In B and C, gray lines show results of 500 replicate simulations obtained by bootstrapping 5 kb windows and re-calculating the relationship between distance to the nearest CGI and minor parent ancestry; colored lines indicate the mean of all replicates in sliding 5 kb windows. The number of simulated chromosomes shown in B was chosen to mimic the amount of data used in our analyses, and is one replicate chosen at random.

One implication is that the reliance on PRDM9 to direct recombination may not only impact reproductive isolation between species directly (as in mice, *45*), but also indirectly. For example, if DMIs tend to occur between neighboring genes (*46*), hybrids between species with PRDM9-independent recombination may experience greater negative selection than species that use PRDM9, because recombination events are more likely to uncouple negatively-interacting alleles. On the other hand, in species with PRDM9-independent recombination, genic regions have higher recombination rates and thus may be more likely to be uncoupled from a deleterious background, potentially providing more opportunities for adaptive introgression. As genomic data accumulates for hybridizing species across the tree of life (*47–53*), the importance of recombination mechanisms for the fate of hybrids can soon be systematically evaluated.

## Acknowledgements

We thank Yaniv Brandvain, Erin Calfee, Graham Coop, Jonathan Pritchard, David Reich, Guy Sella, Sonal Singhal, Matthias Steinrücken and members of the Przeworski, Sella, and Pickrell labs for helpful discussions and/or comments on an early version of the manuscript. We thank the Federal Government of Mexico for permission to collect fish and Gaston Jofre for providing fish pictures. This project was supported by R01 GM83098 grant to MP, NSF DDIG DEB-1405232 to MS, and a Harvard Milton Fund grant to MS. MS was supported by a Hanna H. Gray Fellowship from the Howard Hughes Medical Institute.

## Supplementary Materials

Materials and Methods

Table S1-S5

Figures S1-S25

## References

1. J. A. Coyne, H. A. Orr, Speciation. (Sinaeur Associates, Sunderland, MA, 2004).

2. E. H. Stukenbrock, F. B. Christiansen, T. T. Hansen, J. Y. Dutheil, M. H. Schierup, Fusion of two divergent fungal individuals led to the recent emergence of a unique widespread pathogen species. Proceedings of the National Academy of Sciences of the United States of America 109, 10954–10959 (2012).

3. D. Bachtrog, K. Thornton, A. Clark, P. Andolfatto, Extensive introgression of mitochondrial DNA relative to nuclear genes in the Drosophila yakuba species group. Evolution 60, 292–302 (2006).

4. L. H. Rieseberg, J. Whitton, K. Gardner, Hybrid zones and the genetic architecture of a barrier to gene flow between two sunflower species. Genetics 152, 713–727 (1999).

5. R. Cui et al., Phylogenomics reveals extensive reticulate evolution in Xiphophorus fishes. Evolution 67, 2166–2179 (2013).

6. S. Sankararaman et al., The genomic landscape of Neanderthal ancestry in present-day humans. Nature 507, 354–357 (2014).

7. F. Jacobsen, K. E. Omland, Increasing evidence of the role of gene flow in animal evolution: hybrid speciation in the yellow-rumped warbler complex. Molecular Ecology 20, 2236–2239 (2011).

8. J. W. Poelstra et al., The genomic landscape underlying phenotypic integrity in the face of gene flow in crows. Science 344, 1410–1414 (2014).

9. B. M. Fitzpatrick, H. B. Shaffer, Hybrid vigor between native and introduced salamanders raises new challenges for conservation. Proceedings of the National Academy of Sciences of the United States of America 104, 15793–15798 (2007).

10. I. Juric, S. Aeschbacher, G. Coop, The Strength of Selection against Neanderthal Introgression. Plos Genetics 12, (2016).

11. M. Schumer, R. F. Cui, D. L. Powell, G. G. Rosenthal, P. Andolfatto, Ancient hybridization and genomic stabilization in a swordtail fish. Molecular Ecology 25, 2661–2679 (2016).

12. J. P. Masly, D. C. Presgraves, High-resolution genome-wide dissection of the two rules of speciation in Drosophila. PLoS Biology 5, 1890–1898 (2007).

13. D. C. Presgraves, A fine-scale genetic analysis of hybrid Incompatibilities in drosophila. Genetics 163, 955–972 (2003).

14. S. Tang, D. C. Presgraves, Evolution of the Drosophila Nuclear Pore Complex Results in Multiple Hybrid Incompatibilities. Science 323, 779–782 (2009).

15. K. Bomblies et al., Autoimmune response as a mechanism for a Dobzhansky-Muller-type incompatibility syndrome in plants. PLoS Biology 5, 1962–1972 (2007).

16. H.-Y. Lee et al., Incompatibility of Nuclear and Mitochondrial Genomes Causes Hybrid Sterility between Two Yeast Species. Cell 135, 1065–1073 (2008).

17. K. Harris, R. Nielsen, The Genetic Cost of Neanderthal Introgression. Genetics 203, 881–891 (2016).

18. L. H. Rieseberg et al., Major ecological transitions in wild sunflowers facilitated by hybridization. Science 301, 1211–1216 (2003).

19. M. E. Arnegard et al., Genetics of ecological divergence during speciation. Nature 511, 307–311 (2014).

20. C. I. Wu, The genic view of the process of speciation. Journal of Evolutionary Biology 14, 851–865 (2001).

21. A. A. Hoffmann, L. H. Rieseberg, Revisiting the Impact of Inversions in Evolution: From Population Genetic Markers to Drivers of Adaptive Shifts and Speciation? Annual Review of Ecology Evolution and Systematics 39, 21–42 (2008).

22. M. A. F. Noor, K. L. Grams, L. A. Bertucci, J. Reiland, Chromosomal inversions and the reproductive isolation of species. Proceedings of the National Academy of Sciences of the United States of America 98, 12084–12088 (2001).

23. M. W. Nachman, B. A. Payseur, Recombination rate variation and speciation: theoretical predictions and empirical results from rabbits and mice. Philosophical Transactions of the Royal Society B-Biological Sciences 367, 409–421 (2012).

24. N. Barton, B. O. Bengtsson, The barrier to genetic exchange between hybridizing populations. Heredity 57, 357–376 (1986).

25. D. Ortiz-Barrientos, J. Engelstadter, L. H. Rieseberg, Recombination Rate Evolution and the Origin of Species. Trends in Ecology & Evolution 31, 226–236 (2016).

26. Y. Brandvain, A. M. Kenney, L. Flagel, G. Coop, A. L. Sweigart, Speciation and Introgression between Mimulus nasutus and Mimulus guttatus. Plos Genetics 10, (2014).

27. A. M. Kenney, A. L. Sweigart, Reproductive isolation and introgression between sympatric Mimulus species. Molecular Ecology 25, 2499–2517 (2016).

28. S. Renaut et al., Genomic islands of divergence are not affected by geography of speciation in sunflowers. Nature Communications 4, (2013).

29. M. Schumer et al., High-resolution Mapping Reveals Hundreds of Genetic Incompatibilities in Hybridizing Fish Species. eLife 3, (2014).

30. H. S. Fisher, B. B. M. Wong, G. G. Rosenthal, Alteration of the chemical environment disrupts communication in a freshwater fish. Proceedings of the Royal Society London Series B 273, 1187–1193 (2006).

31. M. Schumer, Y. Brandvain, Determining epistatic selection in admixed populations. Molecular Ecology 25, 2577–2591 (2016).

32. P. Andolfatto et al., Multiplexed shotgun genotyping for rapid and efficient genetic mapping. Genome Research 21, 610–617 (2011).

33. M. Schumer, R. Cui, G. Rosenthal, P. Andolfatto, simMSG: an experimental design tool for high-throughput genotyping of hybrids. Molecular Ecology Resources 16, (2015).

34. M. Schumer et al., Assortative mating and persistent reproductive isolation in hybrids. Proceedings of the National Academy of Sciences of the United States of America 114 10936–10941 (2017).

35. Z. Baker et al., Repeated losses of PRDM9-directed recombination despite the conservation of PRDM9 across vertebrates. eLife 6, (2017).

36. R. Do et al., No evidence that selection has been less effective at removing deleterious mutations in Europeans than in Africans. Nature Genetics 47, 126–131 (2015).

37. Y. B. Simons, G. Sella, The impact of recent population history on the deleterious mutation load in humans and close evolutionary relatives. Current Opinion in Genetics & Development 41, 150–158 (2016).

38. B. Vernot, J. M. Akey, Resurrecting Surviving Neandertal Lineages from Modern Human Genomes. Science 343, 1017–1021 (2014).

39. K. Prufer et al., The complete genome sequence of a Neanderthal from the Altai Mountains. Nature 505, 43–49 (2014).

40. D. Enard, D. Petrov, RNA viruses drove adaptive introgressions between Neanderthals and modern humans. bioRxiv, (2017).

41. F. Racimo, S. Sankararaman, R. Nielsen, E. Huerta-Sanchez, Evidence for archaic adaptive introgression in humans. Nature Reviews Genetics 16, 359–371 (2015).

42. S. Sankararaman, S. Mallick, N. Patterson, D. Reich, The Combined Landscape of Denisovan and Neanderthal Ancestry in Present-Day Humans. Current Biology 26, 1241–1247 (2016).

43. H. A. Orr, The population genetics of speciation - the evolution of hybrid incompatibilities. Genetics 139, 1805–1813 (1995).

44. H. A. Orr, M. Turelli, The evolution of postzygotic isolation: Accumulating Dobzhansky-Muller incompatibilities. Evolution 55, 1085–1094 (2001).

45. B. Davies et al., Re-engineering the zinc fingers of PRDM9 reverses hybrid sterility in mice. Nature 530, 171–176 (2016).

46. Y. Yang, W. Cao, S. Wu, W. Qian, Genetic interaction network as an important determinant of gene order in genome evolution. Molecular Biology and Evolution https://doi.org/10.1093/molbev/msx264, (2017).

47. J. I. Meier et al., Ancient hybridization fuels rapid cichlid fish adaptive radiations. Nature Communications 8, (2017).

48. A. Brelsford, B. Mila, D. E. Irwin, Hybrid origin of Audubon‘s warbler. Molecular Ecology 20, 2380–2389 (2011).

49. B. M. vonHoldt et al., A genome-wide perspective on the evolutionary history of enigmatic wolf-like canids. Genome Research 21, 1294–1305 (2011).

50. J. B. Leducq et al., Speciation driven by hybridization and chromosomal plasticity in a wild yeast. Nature Microbiology 1, (2016).

51. E. B. Taylor et al., Speciation in reverse: morphological and genetic evidence of the collapse of a three-spined stickleback (Gasterosteus aculeatus) species pair. Molecular Ecology 15, 343–355 (2006).

52. L. M. Turner, M. A. White, D. Tautz, B. A. Payseur, Genomic Networks of Hybrid Sterility. PLoS Genet 10, (2014).

53. G. Li, B. W. Davis, E. Eizirik, W. J. Murphy, Phylogenomic evidence for ancient hybridization in the genomes of living cats (Felidae). Genome Research 26, 1–11 (2016).

54. A. Stamatakis, RAxML-VI-HPC: Maximum likelihood-based phylogenetic analyses with thousands of taxa and mixed models. Bioinformatics 22, 2688–2690 (2006).

55. M. Schumer et al., An evaluation of the hybrid speciation hypothesis for Xiphophorus clemenciae based on whole genome sequences. Evolution 67, 1155–1168 (2013).

56. M. Schartl et al., The genome of the platyfish, Xiphophorus maculatus, provides insights into evolutionary adaptation and several complex traits. Nat Genet 45, 567–U150 (2013).

57. M. Malinsky et al., Whole genome sequences of Malawi cichlids reveal multiple radiations interconnected by gene flow. bioRxiv, (2017).

